# On the necessity to include multiple types of evidence when predicting molecular function of proteins

**DOI:** 10.1101/2023.12.18.571875

**Authors:** Valérie de Crécy-Lagard, Manal A. Swairjo

## Abstract

Machine learning-based platforms are currently revolutionizing many fields of molecular biology including structure prediction for monomers or complexes, predicting the consequences of mutations, or predicting the functions of proteins. However, these platforms use training sets based on currently available knowledge and, in essence, are not built to discover novelty. Hence, claims of discovering novel functions for protein families using artificial intelligence should be carefully dissected, as the dangers of overpredictions are real as we show in a detailed analysis of the prediction made by Kim et al ^1^ on the function of the YciO protein in the model organism *Escherichia coli*.

---

Correctly predicting the function of the proteins encoded by whole genome sequences is now a foundational step for most aspects of the biological enterprise. This is far from an easy task and nearly thirty years after the first bacterial genome was sequenced the status of functional annotation of the proteome of most species is still poor^2^. Predicting protein function can be split into two conceptually different problems. The first is to correctly annotate the “knowns” by capturing the existing protein functional knowledge and propagating it to the right set of isofunctional proteins. This process is rife with errors but the most prevalent is the over-annotation of paralogs by propagating annotations too broadly across superfamily subgroups^2^.

The second is to predict the function of the “unknowns,” or of the proteins that have not been linked to a function^3^. Defining the unknown set is not trivial, as definitions of function can vary with the user. Some consider a general annotation such as “transporter” or “ATPase” is sufficient to lose the label of “unknown,” but the prevalent consensus among biocurators is that, for a protein function to be fully known, both a precise molecular function and a biological process must be established, and controlled vocabularies or numbering systems have been established to capture both^4^. By different estimations, the number of “unknowns” in a given genome varies from 20-30% in a handful of model organisms to 50-70% in most sequenced species^5^.

With over 250 million proteins present in the reference database UniProtKB (https://www.uniprot.org/; Release 2023_05 of 08-Nov-2023), predicting protein function cannot be done manually. The last 20 years have seen the incorporation of automation in many steps of the biocuration pipeline^6^. Recently, deep learning-based methods have erupted in the biological fields with successful application in the area of function prediction^7^. Natural language processing and text mining have allowed more efficient filtering of the literature to focus on meaningful papers to curate^8^. One of the exciting developments is the use of transformer base language models that can link protein sequence to Gene Ontologies (GO) describing molecular function, biological process, or localization, or to the EC numbers which are four-part number classifiers describing enzymatic function. These tools can quite accurately propagate the GO information or EC numbers across the protein space, even correctly separating paralogs^9^. As training data improves, reducing bias caused by oversampling of model organisms and commonly known functions, deep-learning could become a key component in the pipelines to correctly annotate the proteins with known functions.

Recently, Kim *et al*. used deep learning with transformer layers (DeepECtransformer) to predict candidate EC numbers for a third of the ∼1500 unknowns present in the model organism *Escherichia coli* K12^1^. The function of three of those YciO, YgfF, and YjdM, were subsequently validated through *in vitro* enzyme activity assays. The authors focused on molecular functions in this study but did not place the predicted functions into the broader biological context.

YciO is a member of the COG0009 family and is a paralog of TsaC/Sua5 (L-threonylcarbamoyladenylate synthase, EC: EC 2.7.7.87), an enzyme that catalyzes the first step in the synthesis of the universal tRNA modification N-6-threonylcarbamoyladenosine or t^6^A. The function of TsaC was first elucidated in 2009^10^ and it was already noted at the time that YciO is a paralog that did not perform the same function *in vivo*, as it did not complement the t^6^A deficient phenotype of a yeast mutant deleted in the gene encoding the TsaC-orthologous enzyme Sua5 (**Fig. 1**), and the *tsaC* gene is essential in *E. coli* even if *yciO* is present in the same genome, and even when the *yciO* gene was overexpressed^11^ (**Fig. 1**). The genetic data therefore show quite convincingly that YciO does not have the same function as TsaC *in vivo*. Gene neighborhood and structural data suggest the actual function of YciO could be related to RNA metabolism; the structure shows a broad positively charged surface predicted to interact with RNA (**Fig. 2**), and in many species *yciO* genes are in operons with *yciV* genes (**Fig. 1**), which encode the recently characterized RNase AM, a 5′ to 3′ exonuclease that matures the 5′ end of all three ribosomal RNAs in *E. coli*^12^.

**Figure 1.**
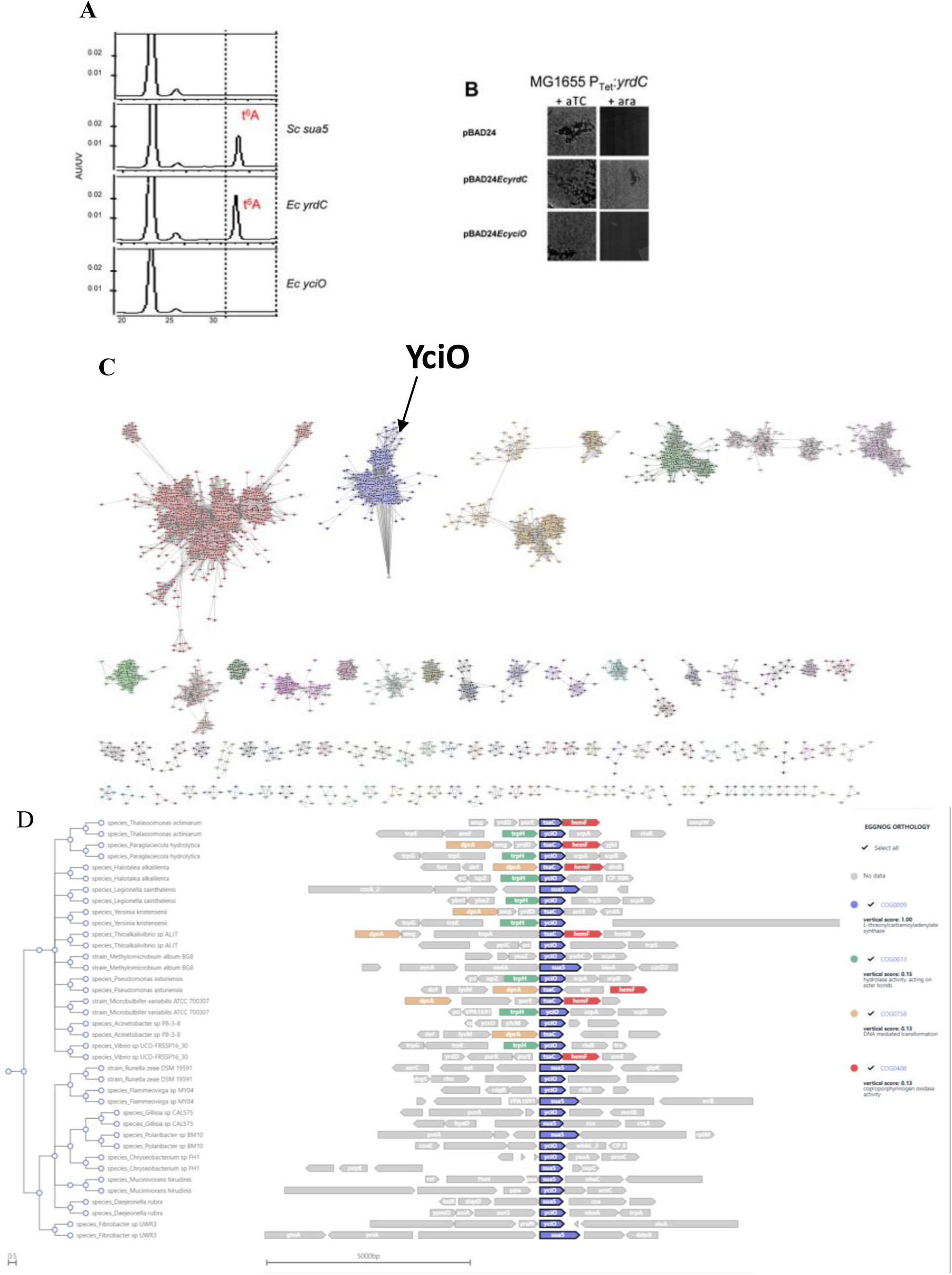
YciO is a paralog of TsaC/Sua5 that does not fulfill the same function *in vivo*. Expression of *E. coli yciO* does not complement t^6^A deficiency of yeast *sua5* deletion (A) nor *tsaC* essentiality in *E. coli* (B). Sequence Similarity Network of the PF01300 family (https://efi.igb.illinois.edu/efi-est/), generated with alignment score threshold of 70, limited to proteins between 200 to 350 aa in size and using an 80% identity RepNode network (all connected sequences that share 85% or more identity are grouped into a single node), and limited to clusters over 10 proteins in size. YciO proteins are only found in one group, all others groups contain predicted or validated TsaC/Sua5 proteins (C). Analysis of COG0009 members automatically generated by Gecoviz (https://gecoviz.cgmlab.org) showing genomes with YciO and TsaC/Sua5 paralogs that show the clustering of *yciO* genes with *yciV*/*trpH* genes (D).

**Figure 2:**
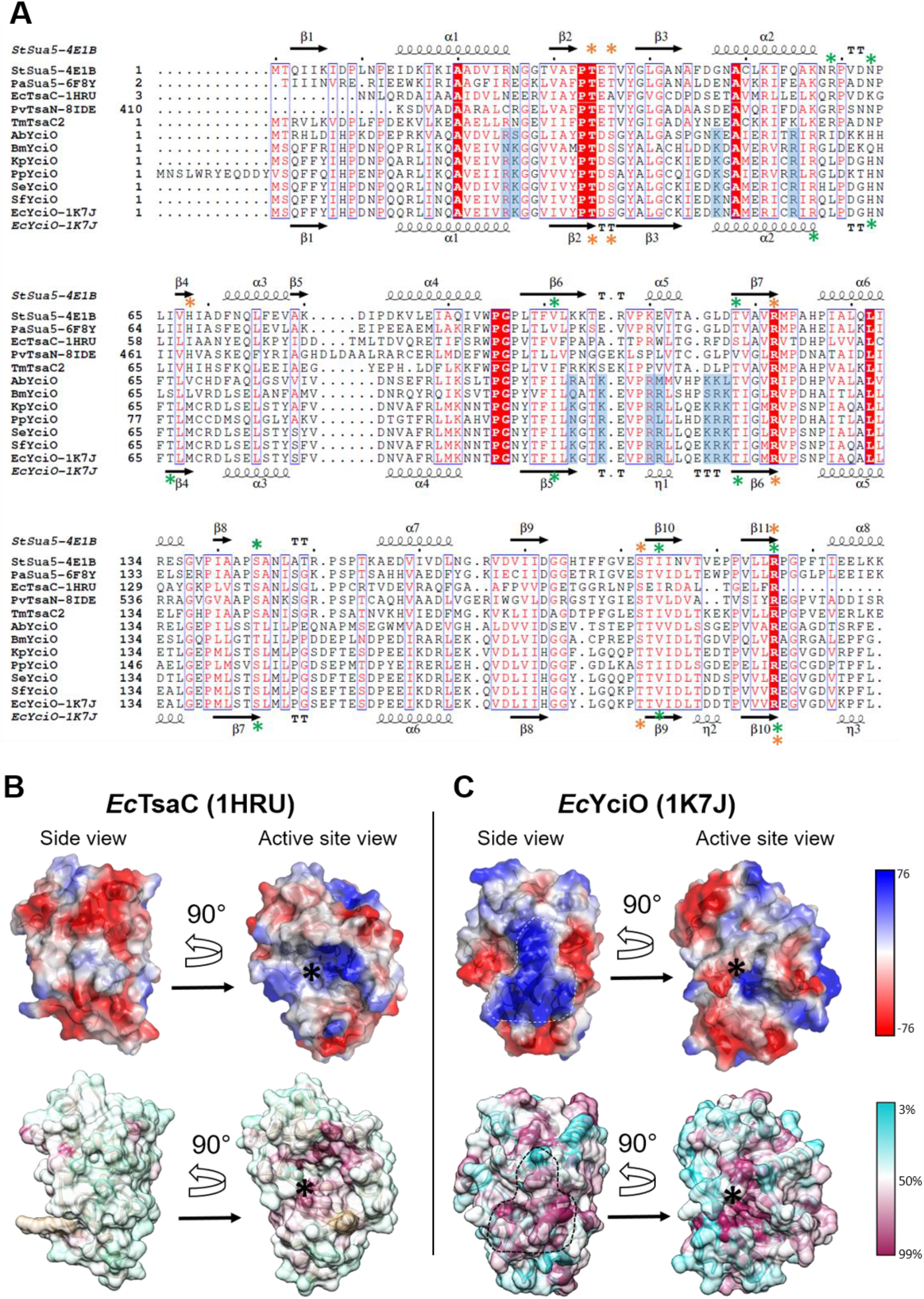
Conserved YciO-specific molecular surface features. **A)** Structure based multi-sequence alignment of TsaC proteins, the TsaC domains of Sua5 proteins and YciO proteins, derived from available crystal structures and AlphaFold structural models, and generated using PROMALS3D (http://prodata.swmed.edu/promals3d/promals3d.php) and Espript (https://espript.ibcp.fr/ESPript/ESPript/). For crystal structures, the PDB IDs are indicated in the sequence name after the hyphen. Secondary structure elements from the crystal structures of *E. coli* TsaC and *E. coli* YciO are displayed above and below the sequences, respectively. YciO-specific conserved basic residues forming the positively charged surface patch of YciO are shaded in blue. Stars above and below the alignment indicate the crystallographically observed substrate binding residues in *St*Sua5, and the corresponding putative substrate binding residues in *Ec*YciO, respectively. Green stars indicate the Mg^2+^-ATP binding residues. Orange stars indicate the binding residues for the _L_-Thr substrate. *Pv*TsaN is the TsaC domain of Pandoravirus TsaN. *Ec*: *Escherichia coli, Pa*: *Pyrococcus abyssi, St*: *Sulfurisphaera tokodaii, Tm*: *Thermotoga maritima* (by the authors, unpublished), *Ab*: *Actinomycetales bacterium, Bm*: *Burkholderia multivorans, Kp*: *Klebsiella pneumoniae, Pp*: *Pseudomonas putida, Se*: *Salmonella enterica, Sf*: *Shigela flexneri*. **B**, **C)** Surface representations of the crystal structures of *Ec*TsaC (**B**) and *Ec*YciO (**C**), color coded by electrostatic potential (top) and positional conservation score (bottom). The color keys for both panels are shown on the right. The conserved YciO-specific positively charged surface patch (6% of total molecular surface area) is encircled with dashed line. The percent conservation score was calculated using the program AL2CO (http://prodata.swmed.edu/al2co/al2co.php), 4000 TsaC/Sua5 sequences and 1500 YciO sequences. The active center (or putative active center for YciO) is marked with an asterisk.

Using the DeepECtransformer platform, Kim *et al*. predicted that YciO had the same function as TsaC, and they presented activity data (albeit missing a no-enzyme control) showing that it could catalyze the same reaction, the synthesis of L-threonylcarbamoyladenylate from ATP, _L_-threonine and CO_2_, *in vitro*. A structure-based multi-sequence alignment comparing YciO and TsaC/Sua5 does show that the active site residues are very similar between the two proteins, consistent with a similar catalytic function (**Fig. 2A**). However, the apparent *in vitro* activity reported by Kim *et al*. must not represent the real biological function, i.e. t^6^A synthesis on tRNA, as integrating the literature (genetic data) and the comparative genomic (physical clustering) data shows that YciO and TsaC/Sua5 must be fulfilling different, but possibly chemically related, biological functions *in vivo*. Consistent with this, comparison of the surface charge distribution of the two protein subgroups reveals that YciO proteins exhibit a highly conserved positively charged surface patch that is absent in TsaC/Sua5 proteins (**Fig. 2B and 2C**), suggesting possible interactions with different partner macromolecules. Notably, the *E. coli* YciO activity reported by Kim *et al*. (0.14 nM/min rate of production of the reaction byproduct PPi) is more than four orders of magnitude weaker than that of *E. coli* TsaC (2.8 μM/min) at the same enzyme concentration^13^ and similar reaction conditions, consistent with the possibility of a missing partner or a different biological substrate. In summary, the functional puzzle is far from being solved for proteins of the YciO subgroup.

As discussed above, deep learning approaches can greatly improve the propagation of EC numbers within isofunctional groups starting with an experimentally validated input ^9^. However, as seen in the Kim *et al*. study ^1^, these pipelines are still prone to the over annotation of paralogs, particularly in the absence of an experimentally validated subgroup member that can provide ground truth for the functional call; they are designed to classify and propagate established knowledge based on training sets and cannot propose novel functional roles not found in the training data. Further improvement of deep learning approaches that considers gene neighborhood information of each protein sequence and the functional roles of neighbors could help with the issue of overpropagation^14^, as could the use of other function descriptors, such as reaction SMILES, from which deep learning methods might learn general rules of chemistry. Introducing phylogenetics, genomic context, structural biology, and experimental findings from free text language as training data remains a challenge for Artificial Intelligence models because the data must be collected, standardized, harmonized, and converted into a machine-learning friendly format. There is no such integrative database available now.

These limitations are not well understood by authors, reviewers, readers, or even biocurators. Therefore, they can lead to swift propagation of incorrect annotations in databases, making them all the more difficult to fix^15^. Some of these mistakes can be easily avoided, namely, by careful literature searches or integration of other types of evidence, such as pathway reconstructions or physical clustering. Had the authors of the Kim *et al*. study performed a PaperBLAST (https://papers.genomics.lbl.gov/cgi-bin/litSearch.cgi) search with the YciO sequence they would have found the 2009 study showing that YciO does not have the same *in vivo* role as TsaC, which would have allowed them to put their EC number prediction and *in vitro* results into a more nuanced perspective.

## Acknowledgments

This work was funded by NIGMS (GM110588) to MAS and VDC. We thank Alan J. Bridge (SIB) and Raquel Diaz (UF) for critical reading and input on the manuscript.

## Contributions

VDC wrote the first draft of the manuscript and did the analyses for Fig. 1. MAS did the analyses for Fig. 2 and edited the manuscript. Both polished the final draft.

